# HPAanalyze: An R Package that Facilitates the Retrieval and Analysis of The Human Protein Atlas Data

**DOI:** 10.1101/355032

**Authors:** Anh Nhat Tran, Alex M. Dussaq, Timothy Kennell, Christopher D. Willey, Anita B. Hjelmeland

**Author notes:** Corresponding authors: Anh Nhat Tran, Anita Hjelmeland, THT 948, 1900 University Blvd, Department of Cell, Developmental and Integrative Biology, University of Alabama at Birmingham, Birmingham, AL 35294. Additional contact information: Alex M. Dussaq, 121 Shelby Biomedical Research Building, University of Alabama at Birmingham Birmingham, AL 35294, Timothy Kennell, 121 Shelby Biomedical Research Building, University of Alabama at Birmingham, Birmingham, AL 35294, Christopher Willey, 176 Facility Building, University of Alabama at Birmingham, Birmingham, AL 35294.

## Abstract

**Background:** The Human Protein Atlas (HPA) aims to map human proteins via multiple technologies including imaging, proteomics and transcriptomics. Access of the HPA data is mainly via web-based interface allowing views of individual proteins, which may not be optimal for data analysis of a gene set, or automatic retrieval of original images.

**Results:** HPAanalyze is an R package for retrieving and performing exploratory analysis of data from HPA. HPAanalyze provides functionality for importing data tables and xml files from HPA, exporting and visualizing data, as well as downloading all staining images of interest. The package is free, open source, and available via Bioconductor and Github.

**Conclusions:** HPAanalyze integrates into the R workflow via the tidyverse philosophy and data structures, and it can be used in combination with Bioconductor packages for easy analysis of HPA data.

## BACKGROUND

The Human Protein Atlas (HPA) is a comprehensive resource for exploration of the human proteome which contains a vast amount of proteomics and transcriptomics data generated from antibody-based tissue micro-array profiling and RNA deep-sequencing (*1–7*). The program has generated protein expression profiles in human non-malignant tissues, cancers, and cell lines with cell type-specific expression patterns via an innovative immunohistochemistry-based approach. These profiles are accompanied by a large collection of high-quality histological staining images that are annotated with clinical data and quantification. The database also includes classification of proteins into both functional classes (such as transcription factors or kinases) and project-related classes (such as candidate genes for cancer). Starting from version 4.0, the HPA includes subcellular localization profiles based on confocal images of immunofluorescently stained cells. Together, these data provide a detailed picture of protein expression in human cells and tissues, facilitating tissue-based diagnostic and research.

Data from the HPA are freely available via proteinatlas.org, allowing scientists to access and incorporate the data into their research. Previously, the R package hpar has been created for fast and easy programmatic access of HPA data (*8*). Here, we introduce HPAanalyze, an R package that aims to simplify exploratory data analysis from those data, as well as provide other functions complementary to *hpar*.

## IMPLEMENTATION

HPAanalyze is an R software package with GPL-3 license that is designed for easy retrieval and exploratory analysis of data from HPA. HPAanalyze allows users to quickly import data tables and xml files from HPA and provides a visual summary of the data. All staining images available in HPA can also be downloaded. Data can be obtained for single proteins or a protein set for pathway analysis.

### The Different HPA Data Formats

The HPA project provides data via two main mechanisms: Full datasets in the form of downloadable compressed Tab-Separated Value (TSV) files are available as well as individual entries in Extensible Markup Language (XML), Resource Description Framework (RDF), and TSV formats. The full downloadable datasets include normal tissue, pathology (cancer), subcellular location, RNA gene, and RNA isoform data. For individual entries, the XML format is the most comprehensive: it provides information on the target protein, antibodies, and a summary of each tissue. Also provided are detailed data from each sample including clinical information, immunohistochemistry (IHC) scoring, and image download links.

### HPAanalyze Overview

HPAanalyze is designed to fulfill 3 main tasks: (i) import, subsetting and export of downloadable datasets; (ii) visualization of downloadable datasets for exploratory analysis; and (iii) facilitation of work with individual XML files (Figure 1). This package aims to serve researchers with little programming experience, while also allowing power users to utilize the imported data as desired.

**Figure 1:**
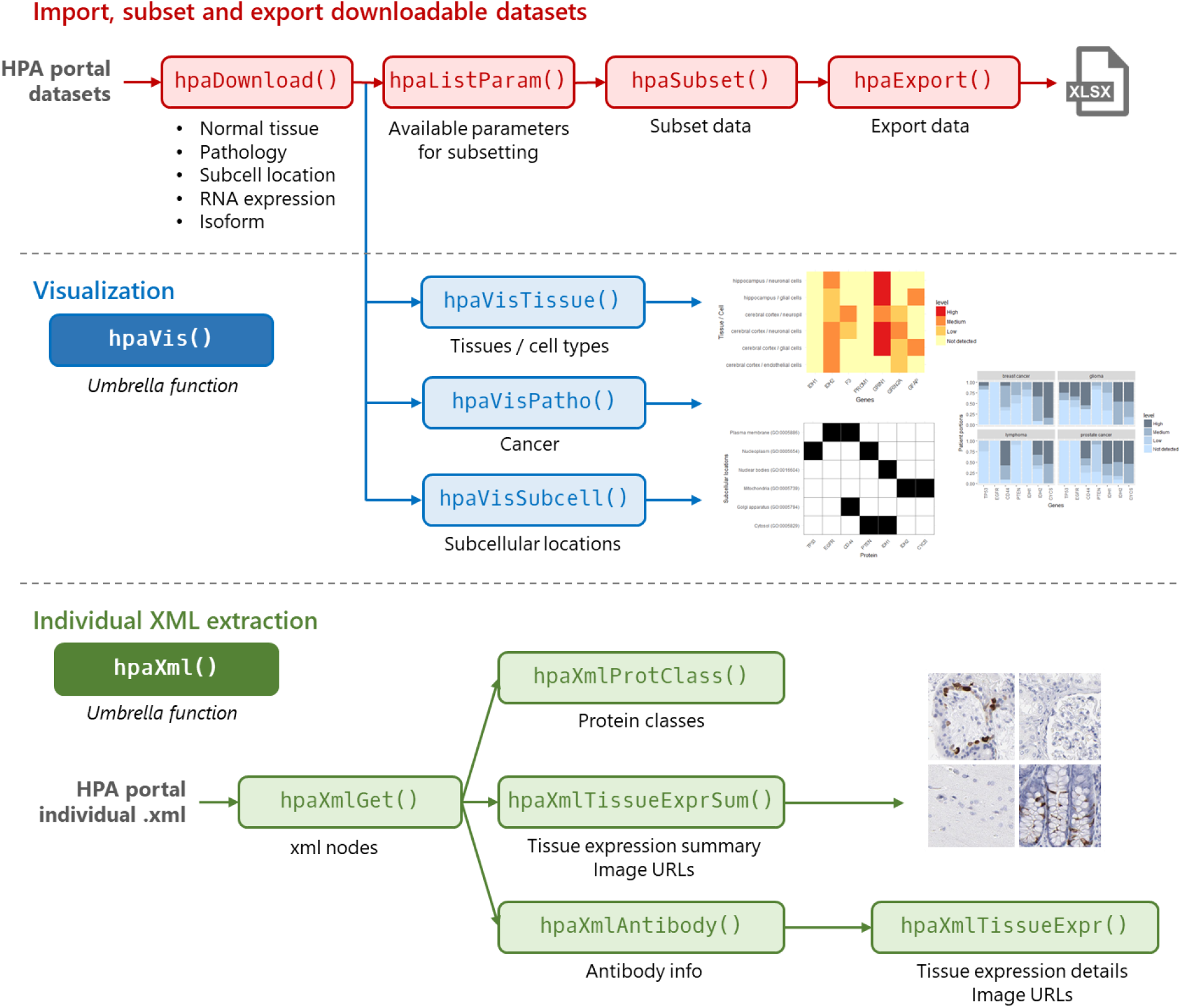
HPAanalyze Workflow. HPAanalyze provides functions for downloading, extracting and visualizing data from HPA. The functions are divided into three different families: 1) hpaDownload for downloadable datasets; 2) hpaVis for quick and customizable visualization; and (3) hpaXml for extracting information from individual XML files.

### Obtaining HPAanalyze

The stable version of HPAanalyze is available via Bioconductor and can be installed with the following code:

~~~
if (!requireNamespace(“BiocManager”, quietly = TRUE))
    install.packages(“BiocManager”)
BiocManager::install(“HPAanalyze”)
~~~

The development version of HPAanalyze is available on Github can be installed with the following code:

~~~
if (!requireNamespace(“devtools”, quietly = TRUE))
    install.packages(“devtools”)
devtools::install_github(“trannhatanh89/HPAanalyze”)
~~~

### Full Dataset Import, Subsetting and Export

The hpaDownload() function downloads full datasets from HPA and imports them into R as a list of tibbles, the standard object of tidyverse. Tibbles can subsequently be subset with hpaSubset() and exported into.xmlx files with hpaExport(). The standard object allows the imported data to be further processed in a traditional R workflow. The ability to quickly subset and export data gives researchers the option to use other non-R downstream tools, such as GraphPad for creating publication-quality graphics, or share a subset of data containing only proteins of interest.

### Visualization

With the intent to aid exploratory analysis, the hpaVis function family takes the output of hpaDownload() [or hpaSubset()] and provides quick visualization of the data. Nevertheless, the standard ggplot (*9*) object output of these functions gives users the option to further customize the plots for publication. All hpaVis functions share the same syntax for arguments: subsetting, specifying colors, and opting to use custom themes.

The first release of the HPAanalyze package includes three functions: hpaVisTissue() for normal tissue samples, hpaVisPatho() for the pathology/cancer samples, and hpaVisSubcell() for subcellular localization data. All operations of this function family can be easily accessed through the umbrella function hpaVis().

### Individual XML Import and Image Downloading

The hpaXml function family imports and extracts data from individual XML entries from HPA. The hpaXmlGet() function downloads and imports data as an “xml_document”/”xml_node” object, which can subsequently be processed by other hpaXml functions. The XML format from HPA contains a wealth of information that may not be covered by this package. However, users can extract any data of interest from the imported XML file using the xml2 package.

In the first release, HPAanalyze includes four functions for data extraction from HPA XML files: hpaXmlProtClass() for protein class information, hpaTissueExprSum() for summary of protein expression in tissue, hpaXmlAntibody() for a list of antibodies used to stain for the protein of interest, and hpaTissueExpr() for complete and detailed data from each sample including clinical data and IHC scoring. hpaTissueExprSum() and hpaTissueExpr() provide download links to obtain relevant staining images, with the former function also providing the option to automate the downloading process. Similar to the hpaVis family, all functionalities of this family may also be accessed through the simple umbrella function hpaXml().

### Compatibility with hpar Bioconductor Package

HPAanalyze was designed to be compatible and complementary to other existing software packages. Table 1 shows the different capibilities of HPAanalyze and hpar, a Bioconductor package optimized for fast acquisition of subsets of HPA data.

**Table 1:** Complementary Functionality Between hpar and HPAanalyze.

| Functionality | Hpar | HPAanalyze |
| --- | --- | --- |
| Datasets | Included in package | Download from server or Import from `hpar` |
| Query | Ensembl id | HGNC symbol for datasets, Ensembl id for XML |
| Data version | One stable version | Latest by default, option to download older |
| Release info | Access via functions | Not applicable |
| View relevant browser page | Via `getHPA` function | Not applicable |
| Visualization | Not applicable | Exploratory via `hpaVis` functions |
| XML | Not applicable | Download and import via `hpaXml` functions |
| Histology image | View by loading browser page | Extract links via `hpaXml` functions |

## TYPICAL WORKFLOWS AND SAMPLE CODES

The HPAanalyze package can be loaded with the following code:

~~~
library(HPAanalyze)
~~~

### Working with HPA Downloadable Datasets

Using HPAanalyze, a typical workflow with HPA downloadable datasets consists of the following steps:

1. Download and import data into R with hpaDownload().
2. View available parameters for subsetting with hpaListParam().
3. Subset data with hpaSubset().
4. Optional: Export data with hpaExport (**Error! Reference source not found.**Figure 1).

The following code can be used to download the histology datasets (normal tissue, pathology, and subcellular location).

~~~
data <- hpaDownload(downloadList = ‘histology’,
                 version = ‘example’)
~~~

The output of the code shows that data can be subset by normal tissue types, normal cell types, cancer types, and subcellular location. The normal_tissue dataset contains information about protein expression profiles in human tissues based on IHC staining. The pathology dataset contains information about protein expression profiles in human tumor tissue based on IHC staining. The subcellular_location dataset contains information about subcellular localization of proteins based on immunofluorescence (IF) staining of normal cells.hpaListParam() function prints a list of available parameters that can be used to subset the downloaded datasets. Below are the first three items in each group:

~~~
hpaListParam(data)
#> $normal_tissue
#> [1] “adrenal gland” “appendix” “bone marrow”
#>
#> $normal_cell
#> [1] “glandular cells” “lymphoid tissue” “hematopoietic cells”
#>
#> $cancer
#> [1] “breast cancer” “carcinoid” “cervical cancer”
#>
#> $subcellular_location
#> [1] “Cytosol” “Mitochondria” “Aggresome”
~~~

Based on the information, the downloaded data may be subset based on genes, tissues, cells and subcellular locations of interest. As an example, the following code filters the datasets for MKI67 (Ki67), breast tissue, and breast cancer.

~~~
hpaSubset(data,
     targetGene = ‘MKI67’,
     targetTissue = ‘breast’,
     targetCancer = ‘breast cancer’) -> ki67_data
~~~

The results (below) showed that Ki67 is expressed at non-detectable-to-medium levels in normal breast tissue, but medium-to-high levels in breast cancer. The data also indicated, with high reliability, that Ki67 is expressed at high levels in the nuclear bodies, nucleoli and nucleus. These data are congruent with the expected pattern of Ki67 expression based on the literature and validate our approach.

~~~
#> $normal_tissue
#> ensembl gene tissue cell_type level reliability
#> <chr> <chr> <chr> <chr> <chr> <chr>
#> 1 ENSG00000148773 MKI67 breast adipocytes Not detected Enhanced
#> 2 ENSG00000148773 MKI67 breast glandular cells Medium Enhanced
#> 3 ENSG00000148773 MKI67 breast myoepithelial cells Not detected Enhanced
#>
#> $pathology
#> ensembl gene cancer high medium low
#> <chr> <chr> <chr> <int> <int> <int>
#> 1 ENSG00000148773 MKI67 breast cancer 2 10 0 #>
#> $subcellular_location
#> ensembl gene reliability enhanced
#> <chr> <chr> <chr> <chr>
#> 1 ENSG00000148773 MKI67 Enhanced Nuclear bodies,’Nucleoli,’Nucleus
~~~

We next sought to facilitate the downstream analysis of data using a non-R software as well as the storage of data subsets for reproducible research. To accomplish this goal, the HPAanalyze package included the hpaExport() function. The hpaExport() function exports data into Excel file format, with each sheet for a dataset. As an example, the code to export the above Ki67 data and generate an.xlsx file called ‘ki67.xlsx’ is as noted below.

~~~
hpaExport(data = ki67_data,
     fileName = ‘ki67.xlsx’)
~~~

### Visualization with The hpaVis Function Family

With the goal of aiding exploratory analysis of a group of target proteins, HPAanalyze provides the ability to quickly visualize data from downloaded HPA datasets with the hpaVis function family (Figure 1). These functions maybe particularly useful for gaining insights into pathways or gene signatures of interest.

The hpaVis functions share a common syntax, where the input is the object generated by hpaDownload() or hpaSubset(). Depending on the function, the target arguments allows the user to choose to visualize vectors of genes, tissue, cell types, etc. All of hpaVis functions generate standard ggplot2 plots, which allow further customization of colors and themes. Currently, the normal_tissue, pathology, and subcellular localization data can be visualized. Additional functions may be added in future releases.

hpaVisTissue() generates a “heatmap”, in which the expression of proteins of interest as measured with quantified IHC staining is plotted for each cell type of each tissue (Figure **2**A). hpaVisPatho() generates an array of column graphs showing the expression of proteins of interest in each cancer (Figure 2B). hpaVisSubcell() generates a tile chart showing the subcellular localization (approved and supported) of proteins of interest (Figure **2**C).

**Figure 2:**
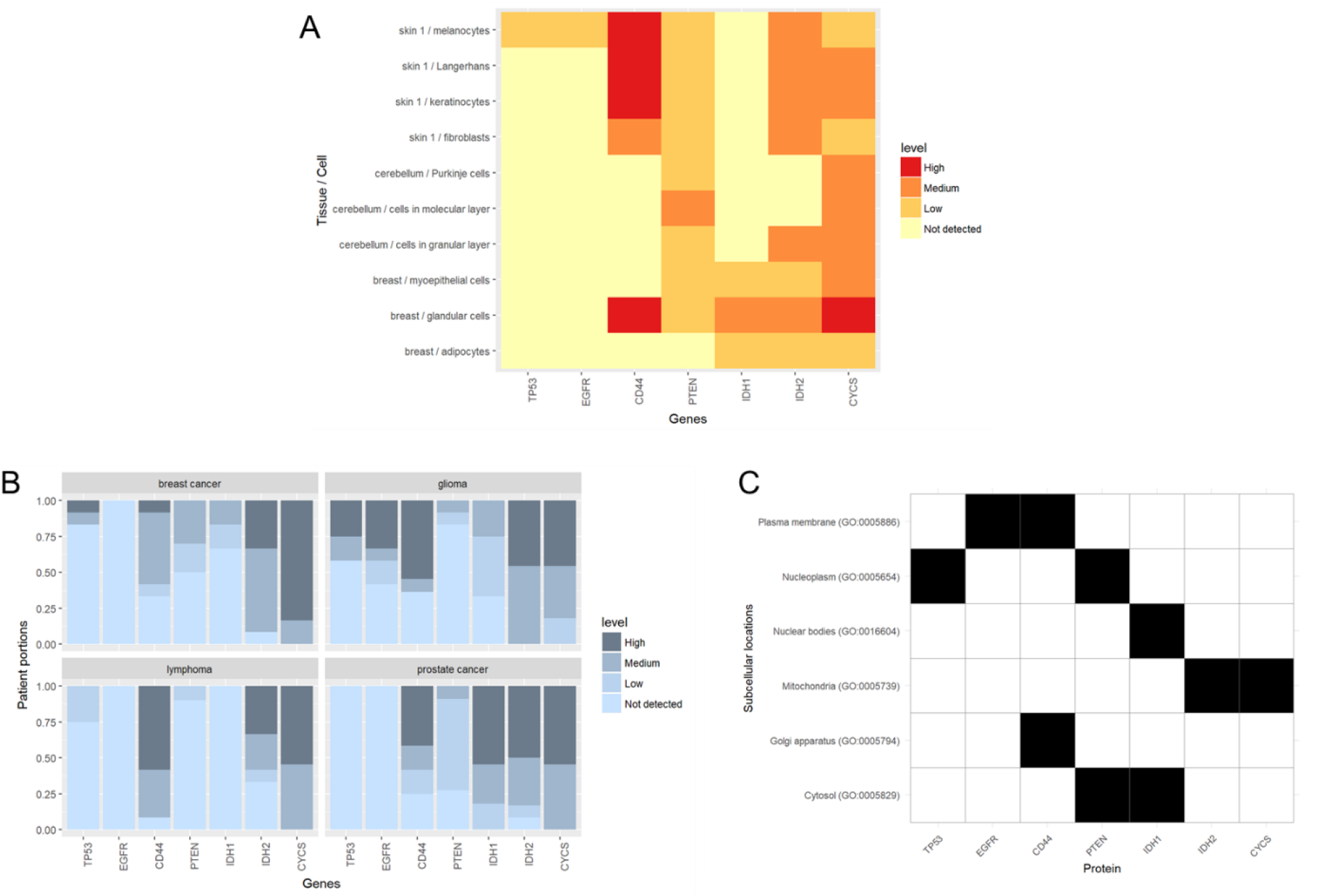
Example of hpaVis Function Family. (A) hpaVisTissue() generates a heatmap of protein expression levels in tissues and cells of interest. (B) hpaVisPatho() creates multiple bar graphs of protein expression in individual target cancers. (C) hpaVisSubcell() visualizes subcellular localization of proteins of interest in a tile chart. Proteins of interest shown via gene name include those implicated in cancer of known subcellular localization: patterns of expression were visualized for Tumor Protein p53 (TP53), Epidermal Growth Factor Receptor (EGFR), Cluster of Differentiation 44 (CD44), Phosphatase and Tensin Homolog (PTEN), Isocitrate Dehydrogenase 1 (IDH1), Isocitrate Dehydrogenase 2 (IDH2), and Cytochrome C (CYCS).

### Working with Individual XML Files for Each Target Protein

The hpaXml function family supports importing and extracting data from individual XML files provided by HPA for each protein. A typical workflow for use of XML files includes the following steps:

1. Download and import XML file with hpaXmlGet().
2. Extract the desired information with other hpaXml functions.
3. Download images of histological stains as currently supported by the hpaXmlTissurExpr() and hpaXmlTissueExprSum() functions (Figure 1).

The hpaXmlGet() function takes a Ensembl gene id (start with *ENSG*) and imports the perspective XML file into R. This function calls the xml2::read_xml() under the hood, hence the resulting object may be processed further with functions from the xml2 package if desired. The protein class of a queried protein can be extracted from the imported XML with hpaXmlProtClass(). The function hpaXmlTissueExprSum() extracts the summary of expression of a protein of interest in normal tissue. The output of this function is 1) a string containing a one-sentence summary, and 2) a dataframe of all tissues in which the protein was positively stained and images of those tissues.

The XML files are the only format of HPA programmatically accessible data that contains information about each antibody and each tissue sample used in the project. hpaXmlAntibody() extracts the antibody information and returns a tibble with one row for each antibody. hpaXmlTissueExpr() extracts information about all samples for each antibody above and returns a list of tibbles. If an antibody has not been used for IHC staining, the returned tibble will be empty. Each tibble contain clinical data (patientid, age, sex), tissue information (snomedCode, tissueDescription), staining results (staining, intensity, location) and one imageUrl for each sample.

## RESULTS/CASE STUDIES

To demonstrate the potential uses of HPAanalyze, we performed example case studies. Each case study was chosen based on the availability of a body of literature that could be used to validate functionality by demonstrating how the resulting data might confirm or complement cancer research.

### Case Study 1: Glioma Pathway Alteration at the Protein Level

As further detailed below, previous studies elucidating pathway alterations in glioblastoma (GBM) have identified frequent amplifications, deletions and mutations of certain genes belonging to Phosphatidylinositol-3-kinase (PI3K)/Mitogen-activated Protein Kinase (MAPK), p53 and Retinoblastoma Protein (Rb) pathways. The level of these proteins in normal brain (hippocampus and cerebral cortex) (Figure **3**A) and glioma (Figure **3**B) was therefore visualized with HPAanalyze. These data were acquired by quantification of IHC stained specimens, so levels of mutant proteins may not be represented. Furthermore, the cancer dataset contains data for all glioma: information about each specific cancer grade is only available via further examination of individual antibodies, which is outside of the scope of this case study.

**Figure 3:**
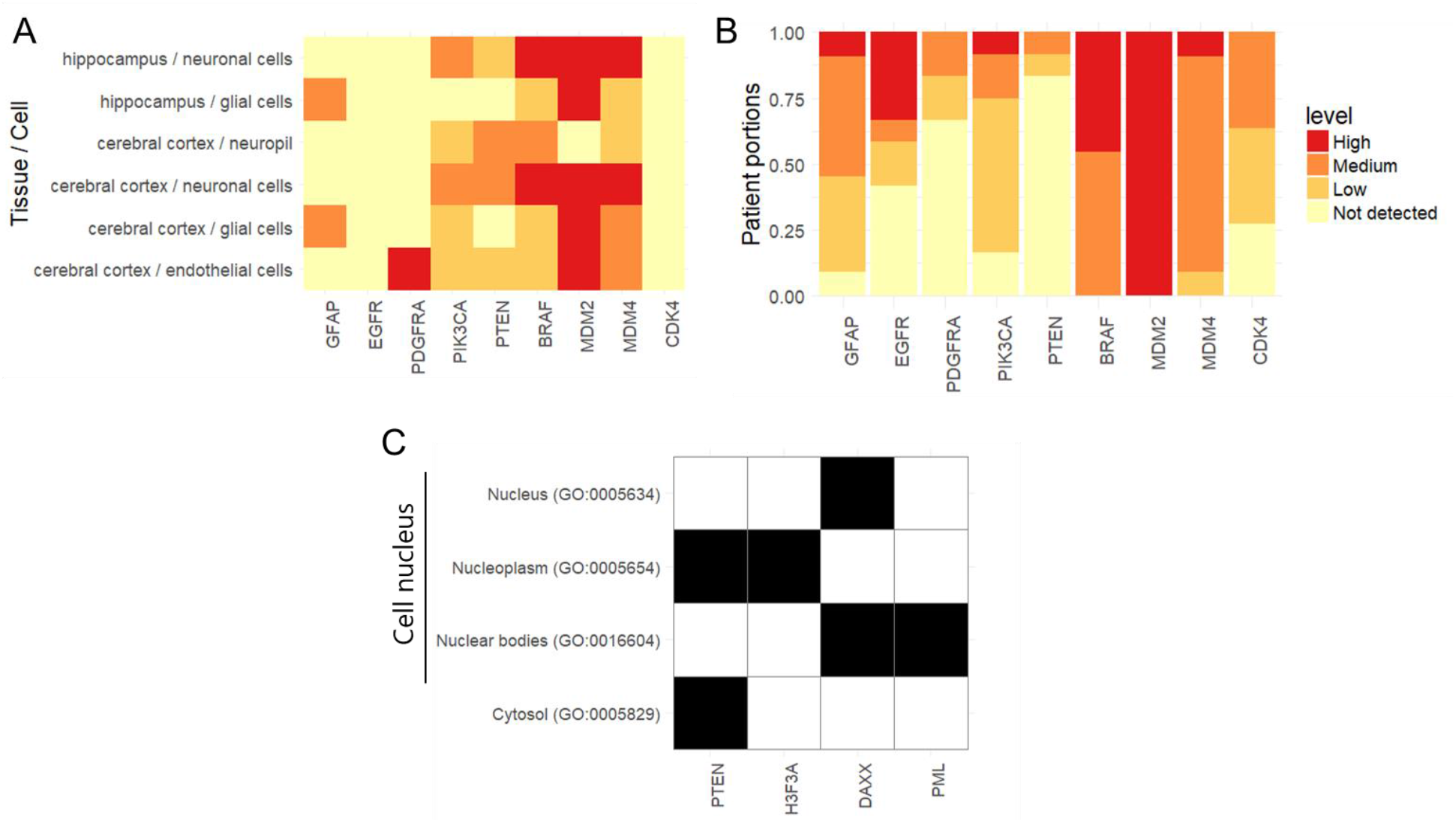
Case Study for Proteins of Interest in Glioblastoma. Expression of proteins frequently altered in GBM in normal (A) or tumor (B) brain tissue. (C) Subcellular localization of proteins associated with non-canonical PTEN function. Molecules of interest shown include Glial Fibrillary Acidic Protein (GFAP), Epidermal Growth Factor Receptor (EGFR), Platelet-derived Growth Factor Receptor Alpha (PDGFRA), Phosphatidylinositol-4,5-Bisphosphate 3-Kinase Catalytic Subunit Alpha (PIK3CA), Phosphatase and Tensin Homolog (PTEN), v-Raf Murine Sarcoma Viral Oncogene Homolog B (BRAF), Mouse Double Minute 2-Like p53 Binding Protein (MDM2), Mouse Double Minute 4 Homolog (MDM4), Cyclin Dependent Kinase 4 (CDK4), H3 Histone Family Member 3A (H3F3A), Death Domain Associated Protein (DAXX), and Promyelocytic Leukemia Protein (PML).

As a positive control, we first evaluated the expression of Glial Fibrillary Acidic Protein (GFAP), a marker for astrocytes/glial cells. In the normal dataset, GFAP expression was found only on glial cells, suggesting that a false positive is unlikely (Figure **3**A). In glioma datasets, GFAP was found to be expressed at mostly medium to high levels (Figure 3B), which is also consistent with the literature.

According to the TCGA data for GBM patients, approximately 90% of patients have alterations in the PI3K/MAPK pathway: Epidermal Growth Factor Receptor (EGFR), Platelet-derived Growth Factor Receptor Alpha (PDGFRA), and PI3K genes are frequently amplified or mutated to gain function (approximately 57%, 10% and 25%, respectively), while Phosphatase and Tensin Homolog (PTEN) is deleted or mutated in 41% of patients (*10*). Data from HPAanalyze supports this pattern, although the proportions are not identical (Figure 3A, 3B). Differences can be attributed to the distinctions between target molecules (DNA/mRNA in TCGA versus protein in HPA) and the number of specimens. One example of the difference can be observed with data regarding v-Raf Murine Sarcoma Viral Oncogene Homolog B (BRAF). BRAF is only amplified/mutated in 2% of TCGA patients, but it is expressed at medium to high levels in all glioma specimens in HPA (Figure 3B).

The p53 pathway is altered in 86% of GBM patients, with amplification of MDM2 (7.6%) and MDM4 (7.2%) leading to the inhibition of p53, which is also highly mutated (*10*). The amplification of MDM2 and MDM4 is reflected at protein levels in HPA: MDM2 is expressed at high levels in all patients and MDM4 at medium levels in most patients (Figure 3B). Similarly, the Rb pathway inhibitor CDK4 was found to be amplified in 14% of patient samples and confirmed by the stark contrast between normal and cancerous samples in HPA (Figure 3A, 3B). The protein CDK4 is not detected in any normal brain cell, while it is present at some level in most glioma samples. These data confirm that HPAanalyze may be useful for comparison of normal and tumor tissue in order to identify or validate molecules of interest with altered expression in cancer.

### Case Study 2: PTEN’s Novel Function through Chromatin-Associated Complexes

PTEN is known as a key tumor suppressor which is frequently mutated in GBM (*10*). Canonically, the protein functions as a phosphatase to dephosphorylate phosphatidylinositol (3,4,5)-trisphosphate (PIP**3**), which leads to inhibition of Akt signaling (*11*). Akt is central to many hallmarks of cancer by promoting cell survival via inhibition of the apoptotic protein Bad, overcoming cell cycle arrest, facilitating glucose metabolism, inhibiting autophagy via regulation of the lysosomal biogenesis controller TFEB, and promoting tumor angiogenesis (*12*).

Since PIP_3_ is a phospholipid that resides on the plasma membrane (*12*), PTEN was once thought to act solely in the cytoplasm. However, a recently published study demonstrated that PTEN also forms complexes with the histone chaperone DAXX and the histone variant H3.3, modulating chromatin association to regulate oncogene expression. This effect is independent of PTEN enzymatic activity (*13*). Congruent with these data, we noted that PTEN was present in both the cytosol and the nucleus (Figure **3**C) in HPA data, suggesting a non-canonical function for PTEN. The subcellular localization of DAXX and H3.3, as well as PML (which interacts with DAXX and regulates PTEN), further corroborate the newly discovered model of PTEN-DAXX-H3.3 gene regulation (Figure 3C).

HPA subcellular localization information for individual proteins is acquired via immunofluorescent staining of human cell lines (*3*). Therefore, the data do not account for various physiological conditions that may relocate proteins nor do the data directly provide evidence of protein-protein interactions. A query of HPA should always be followed by a confirmational study to ensure the validity of the results in any cell type or cancer of interest. Nevertheless, HPAanalyze offers a powerful approach to quickly explore curated and validated antibody-based protein expression data.

### Case Study 3: Protein Expression of GTP cyclohydrolase I (GCH1) pathway

In various cases, the summarized datasets may not be enough to provide a detailed understanding of one protein expression. The HPA program also provided comprehensive data on each individual protein, albeit in the hard-to-parse XML format. HPAanalyze provide functionality that allow XML files to be parsed to extract information on individual specimens and staining quantification for each. We have employed this approach to acquire more details about GCH1 expression in different tumor grades, as well as download IHC images for each sample.

The retrieved data were used in our previously published paper (*14*), which reported the roles of the reactive oxygen species regulator GCH1 in brain tumor (*15*). In short, we found that GCH1 was not expressed in normal brain (cerebellum, cortex, hippocampus or lateral ventricle/caudate) except for in the neuronal cells of the lateral ventricle. However, its expression was detected in glioma samples, and increased with tumor grades. This protein quantification data complemented mRNA expression data in our own study as well as in public datasets such as TCGA and was similar to our IHC staining of patient sample at the University of Alabama at Birmingham.

Furthermore, GCH1 is only one of multiple members of the BH4 biosynthesis pathways. Downstream of GCH1, the de novo pathway also involves 6-pyruvoyltetrahydropterin synthase (PTS) and sepiapterin reductase (SPR), the latter of which has been known to be targeted by multiple well-established sulfa drugs (*16*). BH4 can also be produced via the salvage pathway in which the oxidized product BH2 is converted back to BH4 by dihydrofolate reductase (DHFR) (*17*), a known target for chemotherapy with methotrexate (MTX). While MTX has not proven beneficial for GBM treatment, impacts on GBM BH4 levels are unknown and, if modulated, could provide options for combinatorial approaches.

Our analysis using the HPA datasets showed that, while SPR expression is relatively consistent between normal brain and tumor tissues, other three members of the pathway express at higher levels in glioma tumors than in normal cells (Figure 4**Error! Reference source not found.**A-B). Survival analysis also suggests that, similar to GCH1, the expressions of SPR and DHFR also strongly correlate with worse survival of glioma patients (Figure 4C). Together, the BH4 pathways provide a wide range of targets for drug testing, with readily available small molecule inhibitors.

**Figure 4:**
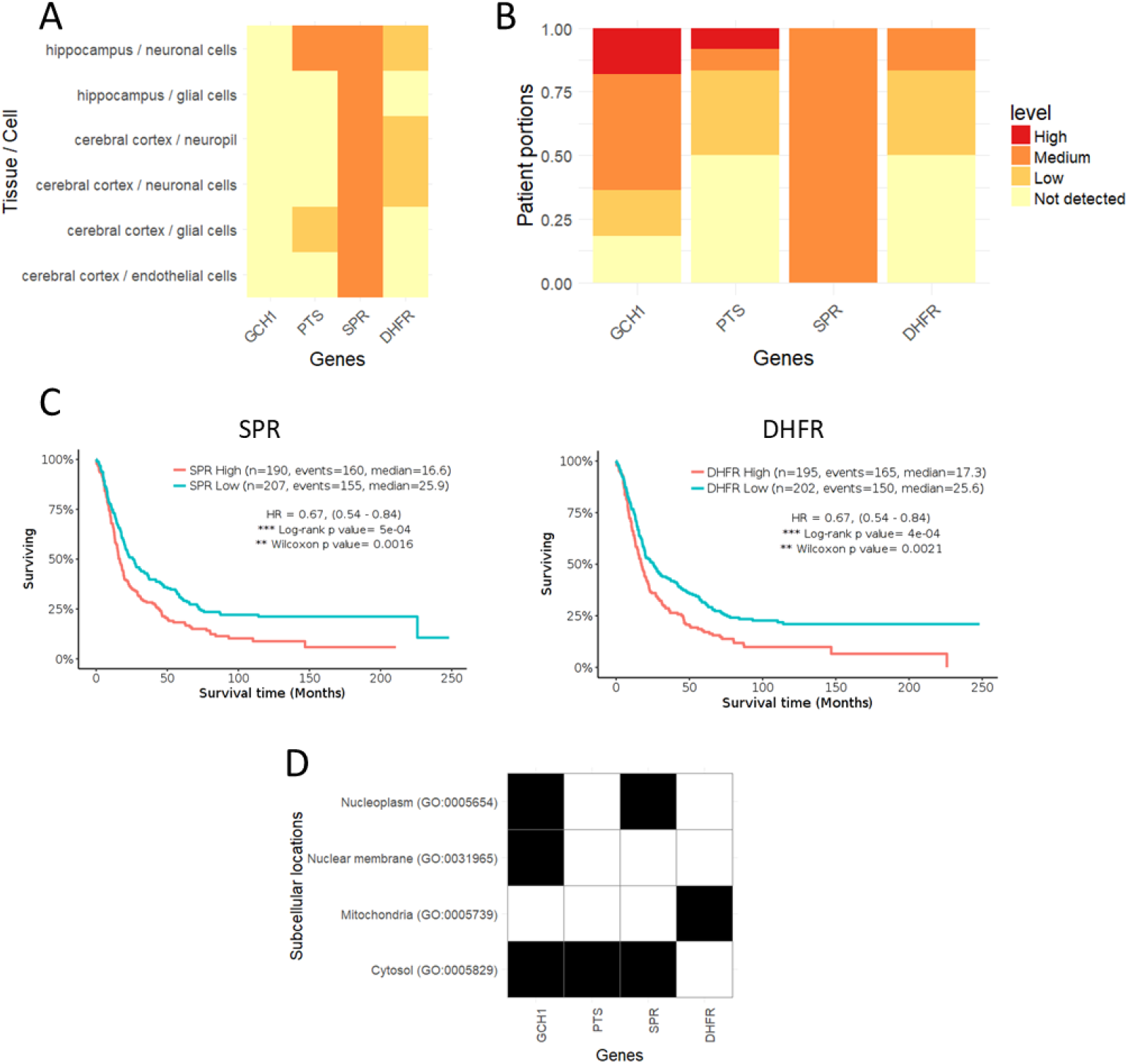
GTP cyclohydrolase I pathway as a potential target for glioma research. (A-B) Protein expression data of genes in the BH4 pathway from the HPA program in (A) normal brain tissue and (B) glioma. (C) Kaplan-Meyer plots showing correlation of SPR and DHFR expression with glioma patient survival, data from the REMBRANDT dataset. (D) Subcellular location data for BH4 pathway members from the HPA program.

Another interesting aspect of BH4 pathway protein expression is the subcellular locations. Based on the data from the HPA program, we found that although all member of the de novo biosynthesis pathway expressed in the cytosol where they function as enzymes to produce BH4 from GTP, the two proteins GCH1 and SPR were also present in the nucleus (Figure 4D). This suggests unknown functions for these proteins, such as transcriptional regulation.

## CONCLUSIONS

Using our R package HPAanalyze, we are able to retrieve, visualize and export data from the HPA program. Although it is a programmatic approach, which requires basic R programming skills, HPAanalyze was built with ease of use and reproducibility in mind, which makes the workflow and syntax very simple and straight-forward. With the case studies, we have also demonstrated how HPAanalyze can be easily integrated into different areas of research to identify new targets or provide more evidence for a working hypothesis. This software package is highly supportive of our research, and we plan to update it with new features and ensure future compatibility with the HPA program.

## AVAILABILITY AND REQUIREMENTS

- Project name: HPAanalyze
- Project home page: https://github.com/trannhatanh89/HPAanalyze
- Operating system(s): All platforms where R is available, including Windows, Linux, OS X
- Programming language: R
- Other requirements: R 3.5.0 or higher, and the R packages dplyr, XLConnect, ggplot2, readr, tibble, xml2, reshape2, tidyr, stats, utils, and hpar
- License: MIT
- Any restrictions to use by non-academics: Freely available to everyone

## LIST OF ABRREVIATIONS

BRAF: : v-Raf Murine Sarcoma Viral Oncogene Homolog B
CD44: : Cluster of Differentiation 44
CDK4: : Cyclin Dependent Kinase 4
CYCS: : Cytochrome C
DAXX: : Death Domain Associated Protein
DHFR: : Dihydrofolate Reductase
EGFR: : Epidermal Growth Factor Receptor
GBM: : Glioblastoma
GCH1: : GTP Cyclohydrolase I
GFAP: : Glial Fibrillary Acidic Protein
H3F3A: : H3 Histone Family Member 3A
HGNC: : HUGO Gene Nomenclature Committee
HPA: : Human Protein Atlas
IDH1: : Isocitrate Dehydrogenase 1
IDH2: : Isocitrate Dehydrogenase 2
IF: : Immunofluorescence
IHC: : Immunohistochemistry
MAPK: : Mitogen-activated Protein Kinase
MDM2: : Mouse Double Minute 2-Like p53 Binding
MDM4: : Mouse Double Minute 4 Homolog
MTX: : Methotrexate
PDGFRA: : Platelet-derived Growth Factor Receptor Alpha
PI3K: : Phosphatidylinositol-3-kinase
PIK3CA: : Phosphatidylinositol-4,5-Bisphosphate 3-Kinase Catalytic Subunit Alpha
PML: : Promyelocytic Leukemia Protein
PTEN: : Phosphatase and Tensin Homolog
PTS: : 6-pyruvoyltetrahydropterin synthase
Rb: : Retinoblastoma Protein
RDF: : Resource Description Framework
SPR: : Sepiapterin Reductase
TCGA: : The Cancer Genome Atlas
TP53: : Tumor Protein p53
TSV: : Tab-Separated Value
XML: : Extensible Markup Language

## DECLARATIONS

### Ethics approval and consent to participate

Not applicable

### Consent for publication

Not applicable

### Availability of data and materials

All data analyzed during this study are publicly available at https://www.proteinatlas.org. The R package is available at https://github.com/trannhatanh89/HPAanalyze.

### Competing interests

Not applicable

### Funding

We appreciate the support of the National institutes of Health R01 NS104339 and R21 NS096531 funds from the Department of Cell, Developmental and Integrative Biology at the University of Alabama at Birmingham.

### Authors’ contributions

ANT created the R package and was a major contributor in writing the manuscript. AMD, TK, Jr., and CDW assisted with package validation and manuscript revision. ABH supervised the project, assisted with data analysis, and revised the manuscript. All authors read and approved the final manuscript.

## Acknowledgements

Not applicable

## COPYRIGHT

© Anh Tran et al. 2018

